# Multi-dimensional spectral gap optimization of order parameters (SGOOP) through conditional probability factorization

**DOI:** 10.1101/438549

**Authors:** Zachary Smith, Debabrata Pramanik, Sun-Ting Tsai, Pratyush Tiwary

**Affiliations:** Biophysics Program and Institute for Physical Science and Technology, University of Maryland, College Park 20742, USA.; Department of Chemistry and Biochemistry and Institute for Physical Science and Technology, University of Maryland, College Park 20742, USA.; Department of Physics and Institute for Physical Science and Technology, University of Maryland, College Park 20742, USA.

## Abstract

Spectral gap optimization of order parameters (SGOOP) (Tiwary and Berne, Proc. Natl. Acad. Sci. **113** 2839 (2016)) is a method for constructing the reaction coordinate (RC) in molecular systems, especially when they are plagued with hard to sample rare events, given a larger dictionary of order parameters or basis functions, and limited static and dynamic information about the system. In its original formulation, SGOOP is designed to construct a 1-dimensional RC. Here we extend its scope by introducing a simple but powerful extension based on the notion of conditional probability factorization where known features are washed out to learn additional and possibly hidden features of the energy landscape. We show how SGOOP can be used to proceed in a sequential and bottom-up manner to (i) systematically probe the need for extending the dimensionality of the RC, and (ii) if such a need is identified, learn additional coordinates of the RC in a computationally efficient manner. We formulate the method and demonstrate its usefulness through three illustrative examples, including the challenging and important problem of calculating the kinetics of benzene unbinding from the protein T4L99A lysozyme, where we obtain excellent agreement in terms of dissociation pathway and kinetics with other sampling methods and experiments. In this last case, starting from a larger dictionary of fairly generic and arbitrarily chosen 11 order parameters, we demonstrate how to automatically learn a 2-dimensional RC, which we then use in the infrequent metadynamics protocol to obtain 16 independent unbinding trajectories. We believe our method will be a big step in increasing the usefulness of SGOOP in performing intuition-free sampling of complex systems. Finally, we believe that the usefulness of our protocol is amplified by its applicability to not just SGOOP but also other generic methods for constructing the RC.

## I. INTRODUCTION

Finding reaction coordinates (RC) and mechanistic pathways in complex systems and processes is a problem of great theoretical and practical interest for which over the decades numerous theoretical and numerical schemes have been proposed.^1–4^ The problem becomes especially complicated in rare event systems, aptly summarized by Chandler and co-workers in their review as the problem of “throwing ropes over rough mountain passes, in the dark”.^2^ Spectral Gap Optimization of Order Parameters (SGOOP) is one such method to construct a RC as a function of candidate order parameters for a given molecular system.^5,6^ This RC encapsulates the most relevant degrees of freedom in the system whose fluctuations must be enhanced in order to accurately sample the thermodynamics and kinetics of metastable states during biased molecular dynamics (MD) simulations such as meta-dynamics or umbrella sampling.^7^ SGOOP was designed keeping rare event systems in mind, where one progressively improves the quality of the RC through rounds of biased simulations performed using it. SGOOP has been demonstrated to be useful for a range of systems such as small peptides and protein–ligand systems, and falls in the broad family of many such related methods that attempt to learn RC for enhanced sampling from sub-optimally biased simulations, such as Ref. 8 and 9. The reason these methods work is at least two fold: (a) irrespective of system complexity, it has been rigorously demonstrated that there exists an optimal one-dimensional RC, given by the normal direction to the isocommittor surfaces,^10–12^ and (b) for the purpose of enhancing the sampling, there is anecdotal evidence that any RC suffices as long as it has sufficient overlap with the true RC.^7,13^ The condition (b) can be rephrased in terms of the timescale separation between slow and fast modes in the system. Namely, the timescales for any process not captured by the RC must be much faster than the slow processes that the RC does encapsulate. SGOOP screens through various putative RCs attempting to maximize this timescale separation, also called spectral gap. In order to do so it uses a maximum path entropy (or Caliber) model that combines any known static or dynamic information about the system,^6,14,15^ and build transition rate matrices along different putative RCs which then directly yield the spectral gap.

SGOOP constructs a one–dimensional reaction coordinate (RC) as a linear or non-linear combination of pre-selected candidate order parameters, which can be thought of as a set of basis functions using which we are trying to describe our problem. Naturally, by considering sufficiently complex combinations of the order parameters (think neural networks) or by making the order parameters themselves sufficiently complex, it should be possible in principle to construct a one–dimensional RC for any given complex process. However for many biomolecular systems of practical relevance that consist of multiple metastable conformations possibly with numerous interconnecting pathways, it might be more desirable to extend the dimensionality of the RC itself. That is, instead of trying to make the 1-d RC more and more sophisticated, it might be computationally cheaper and also physically more interpretable to add a second or even more components to the RC, while still keeping the final dimensionality of the RC much lower than the space of order parameters considered. These other RC components could serve to lift the degeneracy in the first component, and could directly be interpreted in terms of the different pathways or metastable states that they correspond to. A natural question then is how should one go about finding these extra components. The original SGOOP framework could directly be applied to construct a multi-dimensional RC simply by attempting to construct a transition rate matrix on a multi-dimensional grid. This is not very practical since firstly, the dimensionality of the rate matrices scale as *N*^*d*^ × *N*^*d*^, where *d* is each RC-component. Secondly, SGOOP involves calculating the number of barriers discernible in a projection along a given putative RC. This is trivial in 1-*d* but can become tricky and prone to noise related instabilities in higher dimensions.

Here, inspired by Ref. 16 and 17, we develop a simple but powerful extension to SGOOP that makes it possible to sequentially extend the dimensionality of the RC in a straightforward manner. Our approach also makes it possible to quantify when adding further dimensions to the picture is no longer needed. Each additional component is constructed in such a way that it captures features not discernible in the previous components. In this communication, we first develop the key ideas behind our method, which is in fact more generally applicable than SGOOP (see 16 for an illustrative application in the context of deep learning based RC identification), followed by its specific implementation through SGOOP. We then demonstrate the usefulness of our method through different examples of varying complexity, including with model potentials and dissociation of benzene from T4L99A lysozyme in all-atom resolution including explicit TIP3P water. The last system is a popular yet challenging test-case. Here we start with a dictionary of 11 generic order parameters such as protein-ligand and protein-protein distances, and use our automatically learned two-dimensional RC in an infrequent metadynamics framework ^7,18,19^ to calculate its dissociation rate constant and dominant unbinding pathway way, in excellent agreement with previous studies and experiments.^20–24^ We thus believe our method should be of considerable use to the enhanced sampling and molecular simulation communities.

## II. THEORY

### A. Multi–dimensional reaction coordinates through conditional probability factorization

Many previous strategies have been introduced in the past to solve this challenging problem of systematically learning additional hidden variables. For example, inspired by the Marcus theory of electron transfer, Yang and co-workers introduced a method based on considering the generalized force defined by the gradient of the free energy^25^ with respect to the RC. Later, Noe and co-workers introduced a framework inspired by the variational principle in quantum mechanics which constructs a family of RC components. Here we introduce a new framework based on looking at factorized conditional probabilities. In the next sub-section we elaborate the practical implementation of this framework in the context of SGOOP, but it is valid much more generally. Our starting point is a collection of *d* given candidate order parameters *s* = (*s*_1_, *s*_2_, …, *s*_*d*_), and a trial RC 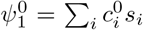. Here the subscript 1 in 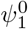 indicates that it is the 1st component of the RC, and the superscript 0 indicates the 0th iteration i.e. starting choice for the same. By following the protocol of Ref. 5 and 26 or any other methods for constructing a 1-d RC, we learn an optimized version of this first component, with different weights {*c*_*i*_}, which we call *ψ*_1_ without any superscripts. Our intention now is to learn a second (and if needed, more) component *ψ*_2_ of the RC that can describe any relevant slow, hidden degrees of freedom, if present, that were not captured by the first component *ψ*_1_. In order to learn *ψ*_2_, we shift our attention from the unbiased or Boltzmann probability distribution *P*_0_ to an auxiliary probability distribution *P*_1_(*s*_1_, …, *s*_*d*_) that is conditional upon what we know about the 1^*st*^ RC component. This distribution thus enhances the features in (*s*_1_, …, *s*_*d*_) space not captured by *ψ*_1_. It is defined by the conditional probability:

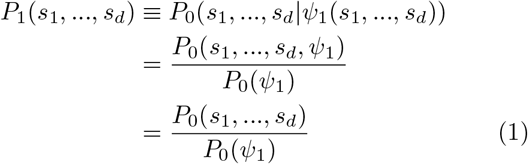

In reaching the last line we have taken into account that given theLvalues of {*s*_*i*_} and a set of coefficients {*c*_*i*_}, *ψ*_1_ = *ψ*_1_(∑_*i*_ *c*_*i*_ *s*_*i*_) is known exactly. Hence the two joint probability distributions in the numerators of the second and third lines of Eq. 1 are equal. Eq. 1 essentially calculates the probability distribution *P*_1_ of the dictionary of order parameters conditional on what we already know about the first component of the RC. In other words, it amounts to sampling the {*s*_1_, *s*_2_, …} space as per Boltzmann distribution *P*_0_, but inverting their weights as per *P*_0_(*ψ*_1_). Given this knowledge, we can now perform a 1-d RC analysis on the probability distribution *P*_1_. If there are no slow, hidden variables, then the probability distribution *P*_1_ should have no additional, orthogonal features on top of what is encapsulated already by the RC *ψ*_1_. We demonstrate this later in Sec. III through numerical examples. However, if there are indeed additional slow variables that were not captured by *ψ*_1_ they will now be expressed through the RC *ψ*_2_ obtained by treating the probability distribution *P*_1_. This reflects the most informative degree of freedom conditional on knowledge of the degrees of freedom captured by *ψ*_1_, and is our second component of the RC. Similar to Eq. 1, the probability distribution for the third component of the RC can be obtained by considering the probability distribution *P*_2_(*s*_1_, …, *s*_*d*_) defined through:

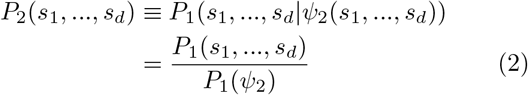

By then repeating this protocol on *P*_*i*-1_ (*s*_1_,…,*s*_*d*_) where *i* ≥ 1 we obtain a sequential set of conditional probability distributions on which we can perform 1-*d* RC optimization for example in the fashion of Ref. 5 and 26: 
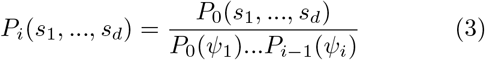

The number of components we choose to identify through this procedure will eventually depend on the problem and sampling method at hand – for example, if the intention is to perform umbrella sampling or metadynamics with the RC, going beyond 2 or 3 components will probably be futile. However for other sampling methods such as Ref. 27 and 28 where in principle one can handle many more biasing variables at the same time, further rounds of the procedure developed here may be applied. A heuristic benchmark for when an additional component *ψ*_*i*+1_ is redundant given the components 1, 2,…, *i* is to examine the correlations between *ψ*_*i*+1_ and *ψ*_1_,…, *ψ*_*i*_ As shown in Sec. III, the usefulness in adding *ψ*_i+1_ can be best judged from examining how correlated or orthogonal are the features in *ψ*_i+1_ to the previous components. Let’s say that one judges that components {*i* + 1,…} do not add any extra information about the slow processes to the representation, and decides to stop the procedure after round *i*. At this point, we can use Eq. 3 to write the full high-dimensional unbiased probability distribution as follows: 
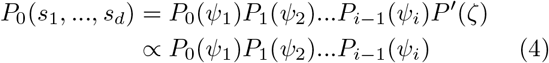
where *P*′(*ζ*) is featureless noise in terms of some more hidden variables *ζ* that we do not care about and thus treat as a constant of proportionality. Thus we have factorized the high-dimensional Boltzmann probability distribution *P*_0_(*s*_1_, *s*_2_,…, *s*_*d*_) as a product of one-dimensional conditional probabilities. This factorization establishes that (i) these variables *ψ*_1_, *ψ*_2_, … and their conditional probability distributions can be learned in a sequential and independent manner as proposed here, and that (ii) these variables together are sufficient to describe the slow modes in the system. We would like to emphasize that Eq. 4 does not imply independence of these variables, i.e. the following is not true: *P*_0_(*s*_1_,…, *s*_*d*_) = *P*_0_(*ψ*_1_)*P*_0_(*ψ*_2_)…*P*_0_(*ψ*_i_). These variables are not independent components and must be treated together.

### B. Multi-dimensional reaction conditional probability factorization through SGOOP

We now describe how the formalism of Sec. IIA can be implemented in practice using SGOOP.^5,6^ Following the notations of Sec. II A, the inputs to SGOOP are a biased simulation performed using a trial RC 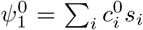, and a short unbiased MD run that gives a time-series of the order parameters *s* = (*s*_1_,*s*_2_,…,*s*_*d*_). Alternatively, the short unbiased MD run could be replaced with estimates of the position-dependent diffusivity tensor.^26^ The biased trajectory is used to obtain estimates of the stationary probability density along various putative RCs, distinguished through values of {*c*_*i*_}, through a postprocessing reweighting procedure^29^, while the unbiased trajectory is used to obtain dynamical constraints needed by the Maximum Caliber framework on which SGOOP is based.^6,14,15^ In SGOOP^5,26^ one spatially discretizes the putative RC *ψ* by defining a grid {*n*} along it, where *n* takes integral values. Let *k*_*mn*_ be the time-independent unbiased rate of transition from grid point *m* to *n* per unit time Δ*t*. Further, let 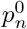 denote the stationary probability of being at any grid point *n* obtained by reweighting the free energy along the respective putative RC^5,6^, and ⟨*N*⟩_0_ represent a dynamic observable which we take here as the average number of first-nearest neighbor transitions in the putative RC grid observed in time Δ*t*. SGOOP^5,26^ uses the following equation to calculate the transition rate for moving from grid point *m* to grid point *n* along any putative RC *ψ*_1_:

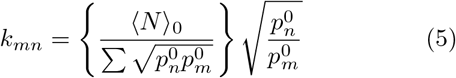

Equivalently, the above equation can be written in a related form by mapping a Smoluchowski equation along *ψ*_1_ term-by-term into a master equation along the same grid, and using diffusion coefficient along *ψ*_1_ instead of MaxCal constraints (see Ref. 26 for details of the mapping). With either form, we can now calculate the eigenvalues of the full transition matrix **K**, where **K**_*nm*_ = −*k*_*mn*_ for *m* ≠ *n* and **K**_*mm*_ = ∑_*m*≠*n*_ *k*_*mn*_. These eigenvalues directly give us the spectral gap for that RC, and by optimizing for the maximal spectral gap by varying the trial coefficients {*c*_*i*_} we learn the best RC, given the static and dynamic information at hand. This optimization can be carried out through a simulated annealing protocol, and gives us an optimized first component of the RC, denoted *ψ*_1_.

So far in this sub-section we have simply summarized SGOOP. This is where we start to extend the protocol. At this point we have an estimate of *ψ*_1_ using which we perform a second metadynamics run biasing *ψ*_1_. Obtaining the probability distribution 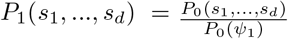 of Eq. 1 from this metadynamics run is embarrassingly trivial: it is the unreweighted, biased probability distribution sampled here. Similar to Eq. 5 we now write down a rate equation along any putative RC *ψ*_2_: 
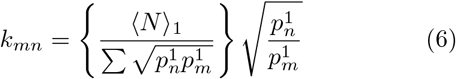

Here 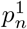 denotes the probability of being at any grid point *n* along a trial 2nd component of the RC, obtained by marginalizing out all other degrees of freedom from *P*_1_(*s*_1_,…,*s*_*d*_). This is a simple binning operation and does not even need the reweighting procedure^29^ for the 1st component, which was needed there to reweight out the effect of biasing along the trial RC 1st component 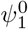. ⟨*N*⟩_1_ represents the average number of first-nearest neighbor transitions in the putative RC grid observed in a time-interval Δ*t* but this time as observed in the simulation performed by biasing *ψ*_1_.

If *ψ*_1_ was truly the only slow degree of freedom, then a search for an RC on the *P*_1_(*s*_1_,…, *s*_*d*_) probability distribution would return no solutions, that is, the optimized RC would be featureless, or even if any features were discovered, they would not be new, and would be already captured by *ψ*_1_ (see Sec. III for examples). However, if indeed a non-trivial RC is found through optimizing the spectral gaps from Eq. 6, we call this as the 2nd component *ψ*_2_ of our RC. We now perform a metadynamics run biasing both *ψ*_1_ and *ψ*_2_, and in principle can repeat this procedure to add as many components as we wish. We re-emphasize that in any successive round, excluding the starting one for *ψ*_1_, there is no need to perform any reweighting, and that the respective biased run itself suffices fully for performing Maximum Caliber based estimates.

## III. RESULTS

Here we first demonstrate our method on two illustrative simple model potentials through a combination of which it can be clearly seen why a second component to the RC might or might not be needed, and how SGOOP can be used to identify the various components. We then apply it to the very challenging test case of benzene dissociation from T4L99A lysozyme in all-atom resolution including explicit TIP3P water, where we are able to accurately simulate the full dissociation process which normally takes hundreds of milliseconds, and calculate the dissociation rate constant *k*_*off*_, a quantity of immense practical relevance in basic biochemistry and drug design^19,30–34^. For the two model systems we considered Eq. 5 in its diffusion constant form as detailed in Ref. 26, and assumed position-independent isotropic diffusivity tensor with no off-diagonal terms. For benzene-T4L99A we considered Eq. 5 directly with MaxCal constraints as detailed in Sec. III B.

### A. Model systems

Both model potentials are represented through sum of three gaussians, and an overall restraining potential, and are in *k*_*B*_*T* units where *k*_*B*_ is Boltzmann’s constant and *T* is the temperature of the system. We use a numerical approach for these two model potentials using analytical/numerical estimates of different reweighted free energies and probability distributions. We did perform MD as well on these potentials (hence the need for restraining potentials), using the full MaxCal version of Eq. 5, and the results were indistinguishable from those reported here.

#### 1. When a 1-component RC is sufficient

The first model potential we considered (Fig. 1) is given by: 
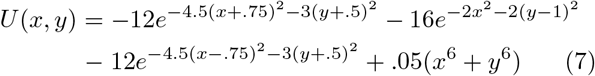

**FIG. 1:**
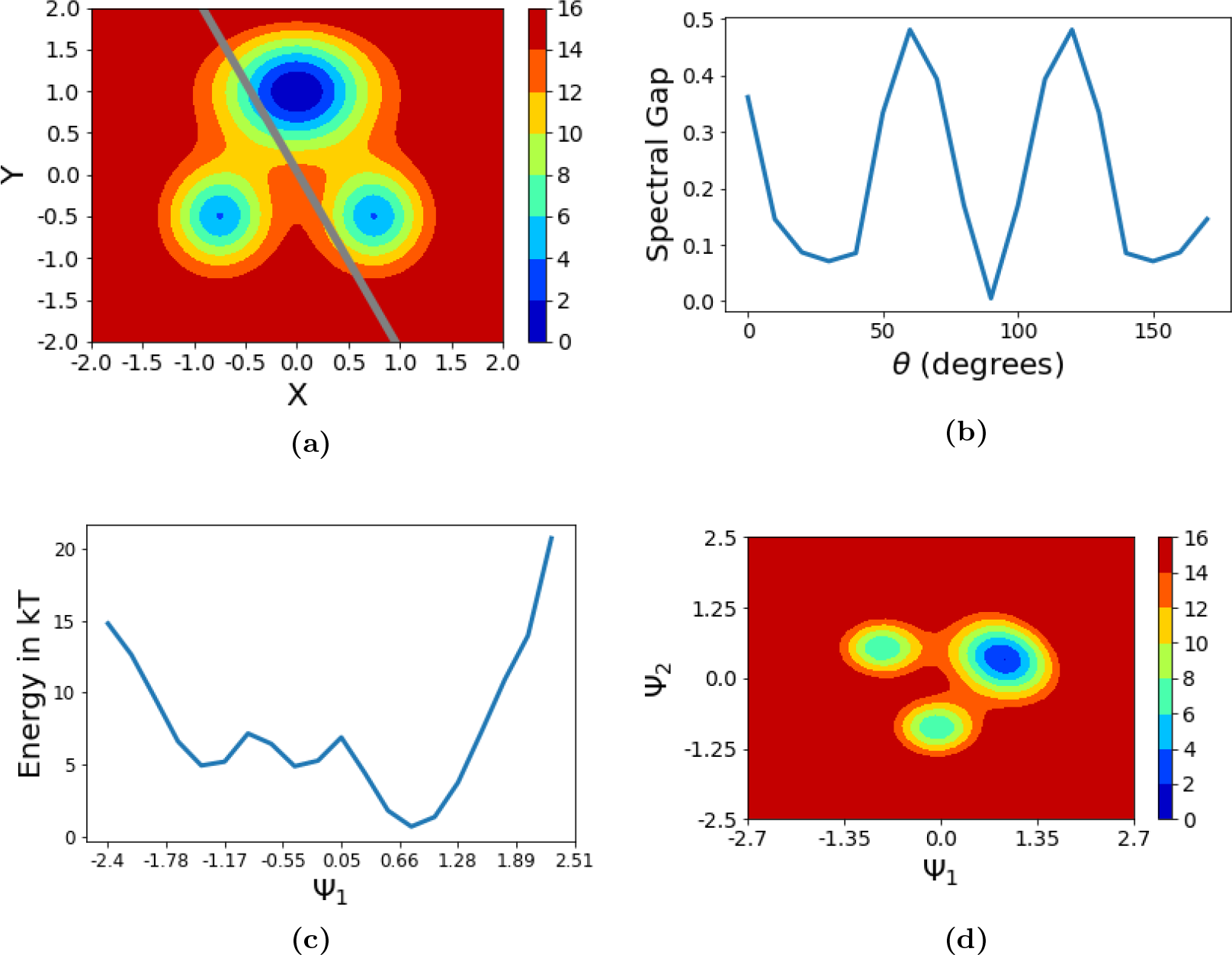
(a) Potential energy contours for Eq. 7. The grey line shows the optimal RC 1st component *ψ*_1_, demarcated through the rotation *θ*=120° measured counter-clockwise from *x*–axis. (b) Spectral gap as a function of *θ* for this potential. (c) Free energy along *ψ*_1_ for this potential. (d) Unbiased free energy (i.e. −*k*_*B*_*T*log *P*_0_(*ψ*_1_, *ψ*_2_)). All energies are in units of *k*_*B*_*T*, while the spectral gap is in arbitrary units since only its relative values concern us.

Here we first identified the first component of the RC, defined as *ψ*_1_(*x*, *y*), as a linear combination of *x* and *y* demarcated through the rotation *θ* measured counterclockwise from *x*-axis. Performing SGOOP here yields spectral gap versus θ profile shown in Fig. 1(b) with two clear maxima at *θ*=60° and *θ*=120°. The two RC solutions are equivalent due to the symmetry of the problem and lead to an identical free energy profile along the RC given in Fig. 1(c). Finally in Fig. 1(d) we have provided the unbiased free energy (i.e. −*k*_*B*_*T*log*P*_0_(*ψ*_1_, *ψ*_1_)), where *ψ*_2_ was calculated to be *θ*=20° by performong SGOOP on *P*_0_(*x*,*y*|*ψ*_1_). We will revisit Fig.1(d) in Sec. III A 2.

#### 2. When a 1-component RC is not sufficient

The second model potential is given by: 
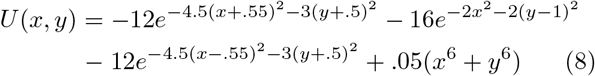

The potential shown in Fig. 2(a) is a modification of the previous potential with the two bottom wells moved closer together as can be seen from Eq. 8. This simple change will cause the overlap between the two wells to be indistinguishable to a single-component linear RC. The spectral gap was optimized as shown in Fig. 2(b) yielding a RC *ψ*_1_ at *θ* = 90° with corresponding unbiased free energy shown in figure Fig. 2(c). *ψ*_1_ was unable to capture all the three energy wells showing that there are hidden degrees of freedom. The conditional probability distribution *P*_1_(*x,y*) = *P*_0_(*x*, *y*|*ψ*_1_(*x*, *y*)) shown in Fig. 2(d) through its associated free energy, was calculated and the spectral gap for the 2nd RC component (Fig. 2(e)) was optimized on this probability distribution. The second component of the RC shown in Fig. 2(d) and given by *θ* = 10° captures a new degree of freedom previously invisible to the first component. Combined these two components can account for both transitions in the *x* and *y* directions despite the *x*–transitions being hidden to *ψ*_1_.

**Fig. 2:**
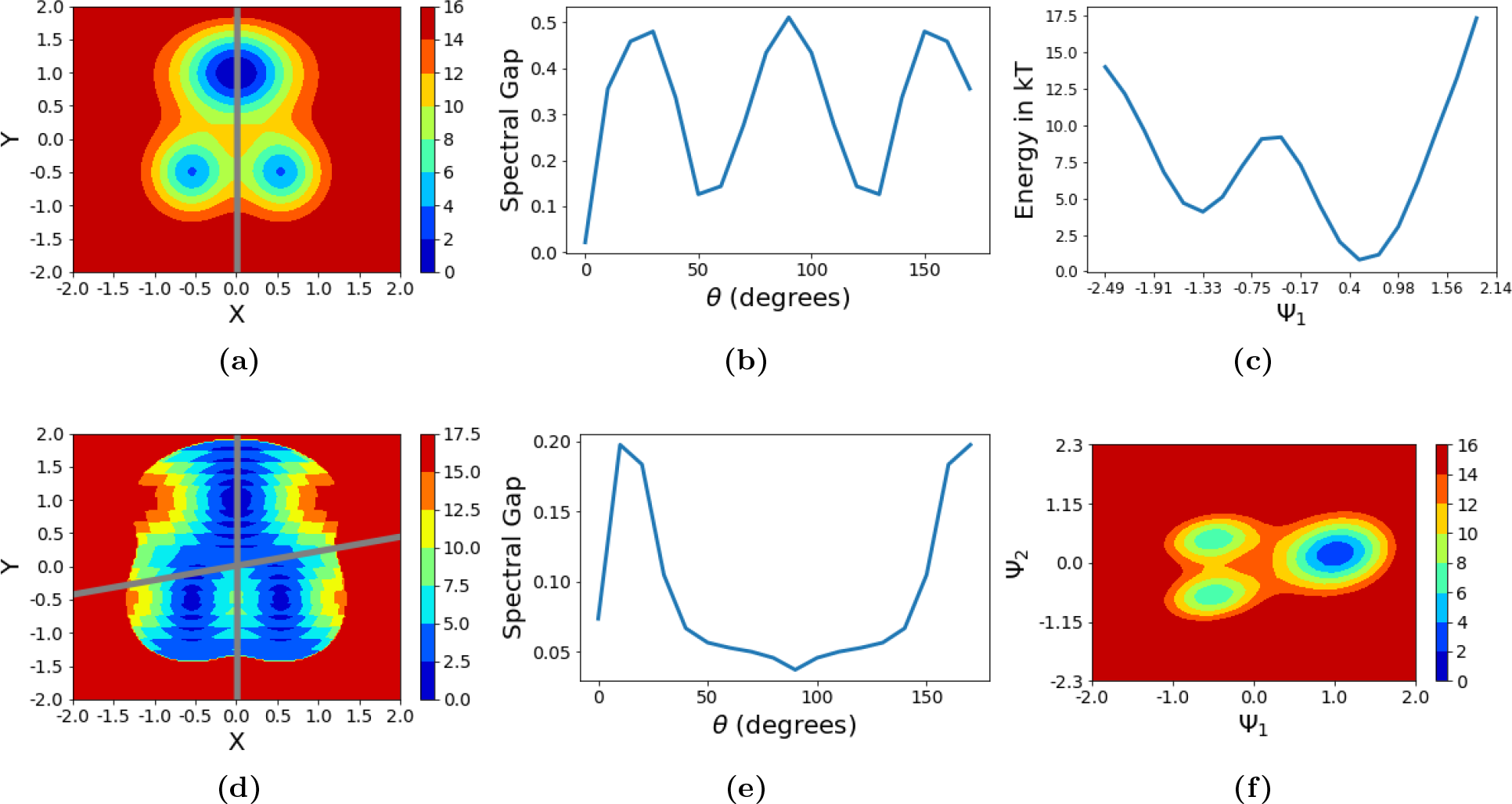
(a) Potential energy contours for Eq. 8. The grey line shows the optimal RC 1st component *ψ*_1_, demarcated through the rotation*θ*=90° measured counter-clockwise from *x*-axis. (b) Spectral gap as a function of *θ* for various *ψ*_1_ choices. (c) Free energy along *ψ*_1_ for this potential. This reaction coordinate only captures the movement between the top well and the bottom wells, and misses the sub-structure of the bottom two wells. (d) Free energy −*k*_*B*_*T*log *P*_1_(*x*, *y*) after conditioning on the estimate of *P*_0_(*ψ*_1_) as per Eq. 1. In addition to the first component to the RC, the second component is also illustrated given by *θ*=10°. (e) Spectral gap as a function of *θ* for *P*_1_(*x*, *y*). (f) Unbiased free energy (i.e. −*k*_*B*_*T*log *P*_0_(*ψ*_1_, *ψ*_2_)). All energies are in units of *k*_B_*T*, while the spectral gap is in arbitrary units since only its relative values concern us.

In Fig. 2(f) we provide the unbiased free energy (i.e. −*k*_*B*_*T*log *P*_0_(*ψ*_1_,*ψ*_2_)) for the potential of Fig. 2(a). This is to be contrasted with the equivalent free energy profile in Fig. 1(d) for the potential of Fig. 1(a). It can be seen from Fig. 2(f) that the stable states demarcated by *ψ*_2_ lie orthogonal to the *ψ*_1_ axis – i.e. they have the same *ψ*_1_ value. However, in Fig. 1(d) this is not the case. The stable states demarcated by *ψ*_2_ can already be distinguished through their *ψ*_1_ values directly. As such, while adding the second component *ψ*_2_ helps in the potential of Fig. 2(a), it does not add any extra information for the potential of Fig. 1(a). This simple heuristic should serve useful in deciding when to add extra components to the RC, as we demonstrate for the next, significantly harder example.

### B. T4 Lysozyme dissociation rate and pathway through infrequent metadynamics

The protocol for extending a RC to multiple components is generally applicable and is expected to be useful for more complex systems for a range of sampling methods. Here we illustrate this through its applicability to infrequent metadynamics, a widely used scheme for recovering unbiased kinetics rates from biased metadynamics simulations.^6,18–21,35–39^ The central idea in infrequentmetadynamics is to perform periodic but infrequent metadynamics is to perform periodic but infrequent biasing of a low-dimensional RC in order to increase the escape probability from metastable states where the system would ordinarily be trapped for extended periods of time. Provided that the chosen RC displays timescale separation and can demarcate all relevant stable states of interest, and if the time interval between biasing events is infrequent compared to the time spent in the transition state (TS) regions, then one increases the likelihood of not adding bias in the TS regions and thereby keeping unbiased the dynamics during barrier crossing itself. This preserves the sequence of transitions between stable states that the unbiased trajectory would have taken. Finally, the acceleration of transition rates through biasing, which directly yields the true unbiased rates, can be calculated through a simple acceleration factor detailed in Ref. 7 and 36. Whether the conditions for the applicability of infrequent metadynamics were met or not can be verified *a posteriori* by checking if the cumulative distribution function for the transition times is Poisson through a Kolmogorov-Smirnoff test developed in Ref. 40. Here one calculates a p-value for the the quality of Poisson fit, and traditionally achieving a value greater than 0.05 is considered safe for reliability.

Using SGOOP we demonstrate how the process of RC selection for infrequent metadynamics can be made almost automatic, starting from a larger dictionary of fairly generic and arbitrarily chosen 11 order parameters. The specific problem considered here is benzene dissociation from the protein T4L99A lysozyme (Fig. 3(a)). This is a well-studied but extremely hard to simulate process due to the debilitating long timescales of milliseconds to seconds, and thus has been studied through different specialized sampling methods. Given the rare event nature of this problem and complex, coupled movements of protein, ligand and even solvent, learning a RC on-the-fly is not a trivial task. We study the process in all-atom resolution using CHARMM22* force field35 for protein, TIP3P water model and CGenFF force field36 for the ligands. Infrequent metadynamics as well has been applied to study this system using exactly the same force-field and MD set-up as we used here. However these and other previous attempts involved putting special effort and fine-tuning into the design of the reaction coordinate to bias during infrequent metadynamics.

**Fig. 3.**
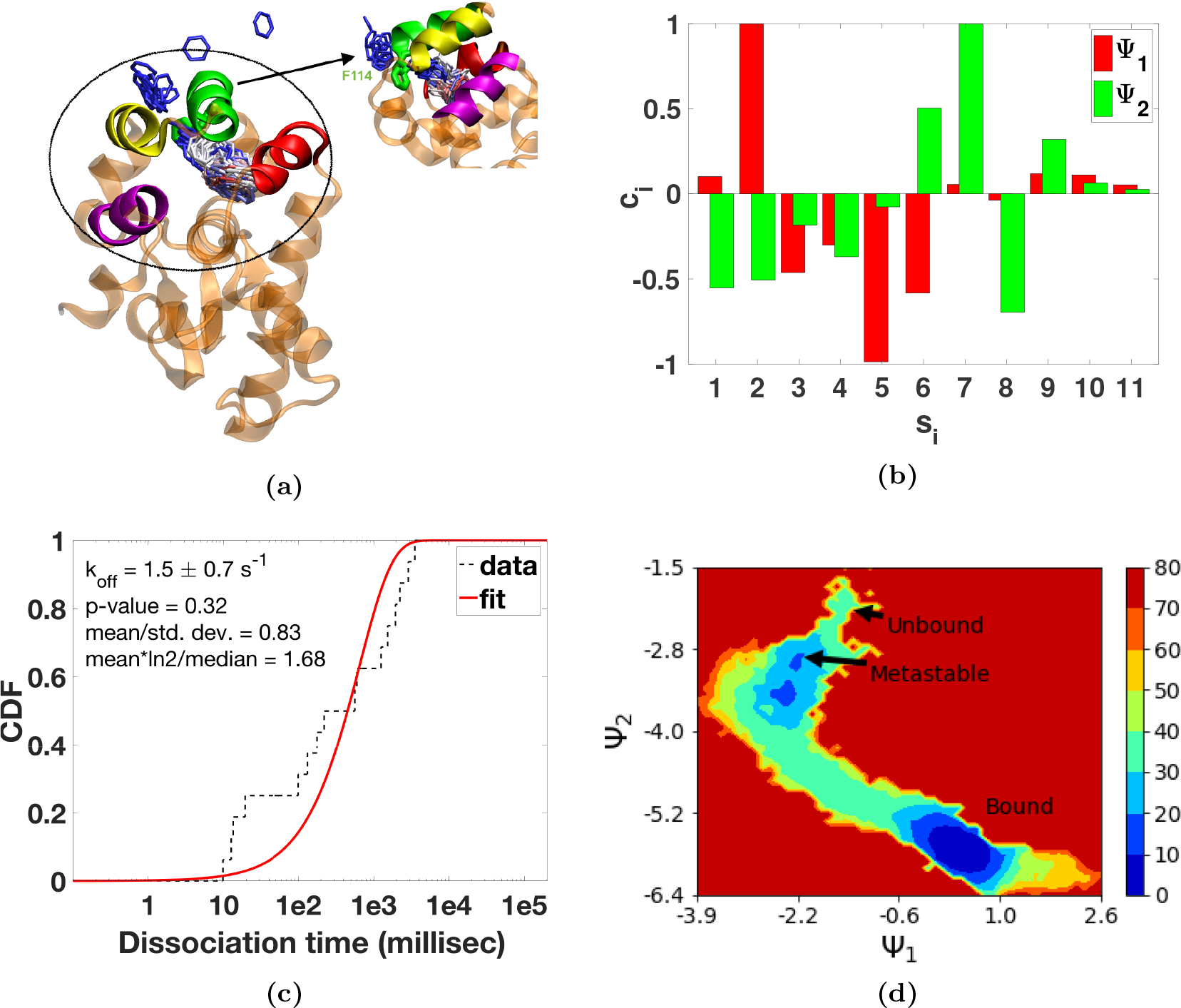
Various details for the benzene-T4L99A system studied here. (a) Secondary structure of the protein, with helices 1, 2, 3 and 4 from Table I shown in colors red, green, yellow and purple respectively. Superimposed is the trajectory of the ligand as it unbinds along the dominant pathway. The trajectory here correspond to 42 *ns* of MD simulation time, with the ligand shown every 10 ps, and colored from red to white to blue as a function of simulation time. 3 clear clusters of states can be seen – bound, metastable and unbound. In the inlay, we have highlighted the residue F114 which acts as a gatekeeper before the ligand reaches the metastable state. (b) A visual depiction of the order parameter weights as tabulated in Table I. (c) Cumulative distribution function (CDF) for the reweighted^18^ unbiased dissociation times (black dashes) obtained from independent infrequent metadynamics simulations, along with a Poisson fit (solid red line). Various statistics indicating the reliable quality of the fit are provided in the inlay. (d) Unbiased free energy (i.e. −*k*_B_*T*log *P*_0_(*ψ*_1_, *ψ*_2_)) in units of *kJ*/*mol*.

Our 11 order parameters {*s*_*i*_} where *i* ranges from 1 to 11 comprise 8 protein-ligand contacts and 3 protein-protein contacts, implemented through simple centre-of-mass to centre-of-mass distances (Table I). Note that we can easily deal with an even higher number of order parameters than 11 without a significant slow-down in the algorithm or the code. This is due to use of simulated annealing protocol for optimization to identify the RC, which is known to scale well with dimensionality.^41^

**Table I:** List of order parameters use to construct RC and their weights in different trial and optimized RC components as learned through SGOOP. Note that no trial values are needed for the second RC-component.

| Order parameter | Type | Definition | Weight $c_i^0$ in trial RC $\psi_1^0 = \sum c_i^0 s_i$ | Weight $c_i$ in optimized $\psi_1 = \sum c_i s_i$ | Weight in optimized $\psi_2 = \sum c_i s_i$ |
| --- | --- | --- | --- | --- | --- |
| $s_1$ | Protein-ligand | Y88CA–ligand | 0 | 0.1019 | -0.5524 |
| $s_2$ | Protein-ligand | A99CA–ligand | 0 | 1 | -0.5068 |
| $s_3$ | Protein-ligand | L133CA–ligand | 1 | -0.4625 | -0.1838 |
| $s_4$ | Protein-ligand | L118CA–ligand | 0 | -0.3011 | -0.3698 |
| $s_5$ | Protein-ligand | V111CA–ligand | 0 | -0.9866 | -0.0767 |
| $s_6$ | Protein-ligand | A130CA–ligand | 0 | -0.5829 | 0.5033 |
| $s_7$ | Protein-ligand | N140CA–ligand | 0 | 0.0548 | 1 |
| $s_8$ | Protein-ligand | A146CA–ligand | 0 | -0.0376 | -0.6962 |
| $s_9$ | Protein-protein | Helix 1 (A82-S90) – Helix 2 (T115-123Q) | 0 | 0.1188 | 0.3192 |
| $s_{10}$ | Protein-protein | Helix 2 (T115-123Q) – Helix 3 (W126-A134) | 0 | 0.1113 | 0.0642 |
| $s_{11}$ | Protein-protein | Helix 3 (W126-A134) – Helix 4 (K147-T155) | 0 | 0.0527 | 0.0260 |

We now provide further details of the implementation as well as results so obtained. We first performed a metadynamics run using trial RC 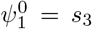. This was done using a relatively frequent and aggressive metadynamics protocol, since the objective was to obtain an approximate estimate of the stationary probability density for use in SGOOP. Specifically we used a well-tempered metadynamics protocol,^29,42^ with initial hill height = 1.5, biasfactor γ = 15, gaussian width σ = .02, and bias added every 1 *ns*. The simulation was performed using GROMACS version 5.1 patch with PLUMED version 2.3.^43,44^ A short unbiased MD run of 18 *ns* was performed in parallel that was used to construct the MaxCal dynamical observable of average number of transitions in any order parameter *s*_*i*_ in 200 *fs*. All simulations were performed in NPT using isotropic Parrinello-Rahman barostat with a time constant of 2 *ps* and modified Berendsen thermostat with a time constant of 0.1 *ps*. From these two runs, we obtained an estimate of the 1st RC component *ψ*_1_ (Table I). To identify the 2nd component we perform metadynamics with similar parameters as for the 1st component, but this time biasing *ψ*_1_ instead of 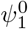. SGOOP is applied to the probability distribution *P*_1_(*s*_1_, *s*_2_,…, s_11_) = *P*_0_(*s*_1_, *s*_2_,…, *s*_11_ |*ψ*_1_). In practice this amounts to ignoring the bias deposited as function of *ψ*_1_ and taking the metadynamics trajectory as is. The same trajectory can also be used for calculating MaxCal constraints. From this we obtain the second component ψ_2_. See Table I and Fig. 3(b) for weights of different order parameters in *ψ*_1_ and *ψ*_2_. In principle we could add further components to the RC. We however note that the objective in this exercise is to perform infrequent metadynamics biasing together the various components of the RC. Since infrequent metadynamics and metadynamics in general becomes extremely slow computationally if one was to use three or more different biasing variables, we stop at this point.

The two components of the RC *ψ*_1_ and *ψ*_2_ are then used in the infrequent metadynamics protocol to construct a two-dimensional bias as a function of these RCs. We perform 16 independent unbinding simulations all starting from the x-ray bound pose with different randomized velocities at 298 K corresponding to Boltzmann distribution. These were performed using a well-tempered metadynamics protocol^29,42^ with initial hill height 1.5 *kJ*, biasfactor γ = 15, gaussian widths 0.1 for both *ψ*_1_ and *ψ*_2_, and bias added every 8 *ps*. Each run is stopped when *s*_1_ reaches 3 *nm*, at which point the ligand is fully solvated and has started freely diffusing. This is our unbound state. From these 16 biased trajectories we find the corresponding unbiased dissociation time estimates through computing the respective acceleration factors, which are in the range of five to seven orders of magnitude. These are then subjected to the Kolmogorov-Smirnov test of Ref. 40, where we obtain a p-value of 0.32, which is well above the recommended and normally used threshold of 0.05, suggesting reliability of the kinetics and associated pathways. Various associated metrics demonstrating quality of Poisson fit are provided in Fig. 3(c). Finally from this fit we obtain a dissociation rate of 1.5±0.7*s*^−1^. This is within error bar agreement with previous infrequent metadynamics using path CVs with the same force-field.^20^ In total our 16 full unbinding trajectories took around 700 *ns* for a process that actually takes a few hundred ms. This reflects the tremendous computational speed-up achieved with respect to unbiased MD, while still recovering unbiased rates well within order of magnitude agreement with results using other methods for an identical force-field parametrization.^20,21^

In Fig. 3(a) we provide an overlaid depiction of a typical dissociation trajectory seen in our simulations, wherein the ligand moves through the crevice between helices 2 and 3 (colored green and yellow respectively). At least 3 distinct clusters can be seen corresponding to the bound state, a metastable unbound state where the ligand is stuck on the surface of the protein, and finally the unbound state when the ligand is freely diffusing in the solvent. In the inlay to Fig. 3(a) we have highlighted the residue F114 which acts as a gatekeeper for the ligand to go from bound state to the metastable state. The exit of the ligand is coupled to the breathing of these helices with respect to each other which opens up space for the ligand to exit. Fig. 3(b) gives the weights of both the RC components, scaled so that every order parameter in Table I ranges between 0 and 1. Note that the unscaled order parameter weights have however been used for Fig. 3(d), since these are what we use while biasing in metadynamics. We would like to highlight that this same pathway has been reported to be the dominant dissociation pathway in other works on this system.^20,22^

It is interesting to examine the weights of the different order parameters in both RC components. The highest weight in *ψ*_1_ is for *s*_2_ which corresponds to ligand separation from the residue A99, which is in the interior of the binding pocket. A roughly equal in magnitude but opposite in sign weight is carried by *s*_5_, reflecting that the ligand moves closer to V111 as it moves away from A99. In *ψ*_2_ the highest weight is for *s*_7_ which corresponds to ligand separation from the residue N140, which lies on one of the two helices surrounding the final ligand exit pathway, and is quite distant from the A99 residue contributing to *s*_2_ which is important for *ψ*_1_. Furthermore, while the protein-protein separation order parameters have non-zero weights in either RC component, these are not the most dominant players. This reflects that the protein breathing motion, while a critical slow process, is not the main driver for dissociation.

Finally, in Fig. 3(d) we provide the unbiased free energy (i.e. −*k*_B_*T*log *P*_0_(*ψ*_1_,*ψ*_2_)) for this system. This is again to be contrasted with the equivalent free energy profiles in Fig. 1(d) and Fig. 2(f) for the potentials of Fig. 1(a) and Fig. 2(a) respectively. It is more similar to the latter: the escape from the barrier between bound and metastable state, as well as out of the metstable state are far better distinguished through their *ψ*_2_ values, and have much more tightly spaced *ψ*_1_ values. As such, here as well adding the second component *ψ*_2_ helps in lifting the degeneracy of states demarcated by *ψ*_1_. Of course, we could have added more components here, but since the objective is to perform infrequent metadynamics, we stopped at two components.

## IV. DISCUSSION

In this work, we have introduced a conditional probability factorization scheme for extending the dimensionality of the reaction coordinate (RC) in a given molecular system. Specifically, here we developed and demonstrated the algorithm in context of the RC optimization method named “Spectral gap optimization of order parameters (SGOOP)”.^5,26^ Our motivation is that often it might be desirable to prefer a multi-dimensional RC with different simple components, over a one-dimensional complex RC. To find such a multi-dimensional RC, the central idea in this work is to progressively “wash out” known features to learn additional and possibly hidden features of the energy landscape. In a sense, this is inspired by the approach of metadynamics^7^ where one gradually builds a memory kernel as a function of a given RC to revisit new parts of the landscape. Here we do an analogous operation in a multi-dimensional RC space, and by building memory of features already learned, we explore additional, hidden low-dimensional features themselves which then can be used to extend the dimensionality of the RC. This higher dimensional RC then gives a more accurate picture of the slow dynamics in the system, and could then also be used to deposit a memory kernel as function thereof. We also want to remark that for the purpose of forming reasonably accurate estimates of the RC {*ψ*_1_, *psi*_2_,…}, we find that poorly converged estimates of the marginal probabilities *P*_0_,*P*_1_ … etc. are already useful, as long as they can be used to at least partially wash out the features already captured. This is line with what has been been reported previously for SGOOP.^5,6^

We demonstrated the usefulness of our approach through three illustrative examples, including the problem of calculating kinetics of benzene unbinding from the protein T4L99A lysozyme. In this last case, we started from a larger dictionary of fairly generic and arbitrarily chosen 11 order parameters, and demonstrated how to automatically learn a 2-dimensional RC, which we then used in the infrequent metadynamics protocol^18,40^ to obtain 16 independent unbinding trajectories. This directly gave us insight into dominant dissociation pathway, as well as the dissociation kinetics.

We believe our method will be a big step in increasing the usefulness of SGOOP in performing intuition-free sampling of complex systems. We also believe that the usefulness of our protocol is amplified by its applicability to not just SGOOP or metadynamics but also other generic methods for constructing the RC and sampling energy landscapes in complex systems. A Python based code implementing the method is available for public use at GitHub.

## ACKNOWLEDGMENTS

We would like to thank Pablo Bravo for coding some of the preliminary parts of SGOOP in Python and João Ribeiro for discussions regarding the central idea in this paper. We thank Deepthought2, MARCC and XSEDE (projects CHE180007P and CHE180027P) for providing the computational resources used to perform this work. ZS would like to thank the Hockmeyer family for financial support during summer of 2018. PT would like to thank University of Maryland Graduate School for financial support through the Research and Scholarship Award (RASA).

## References

1 A. Berezhkovskii and A. Szabo, J. Chem. Phys. 122, 014503 (2005).

2 P. G. Bolhuis, D. Chandler, C. Dellago, and P. L. Geissler, Ann. Rev. Phys. Chem. 53, 291 (2002).

3 M. A. Rohrdanz, W. Zheng, and C. Clementi, Ann. Rev. Phys. Chem. 64, 295 (2013).

4 B. Peters, Reaction rate theory and rare events (Elsevier, 2017).

5 P. Tiwary and B. J. Berne, Proc. Natl. Acad. Sci. 113, 2839 (2016).

6 P. Tiwary and B. J. Berne, J. Chem. Phys. 145, 054113 (2016).

7 O. Valsson, P. Tiwary, and M. Parrinello, Ann. Rev. Phys. Chem. 67, 159 (2016).

8 J. McCarty and M. Parrinello, J. Chem. Phys. 147, 204109 (2017).

9 M. M. Sultan and V. S. Pande, J. Chem. Theor. Comp. 13, 2440 (2017).

10 B. Peters and B. L. Trout, J. Chem. Phys. 125, 054108 (2006).

11 L. Maragliano, A. Fischer, E. Vanden-Eijnden, and G. Ciccotti, J. Chem. Phys. 125, 024106 (2006).

12 S. V. Krivov, J. Chem. Theor. Comp. 9, 135 (2012).

13 P. Tiwary and A. van de Walle, in Multiscale Materials Modeling for Nanomechanics (Springer, 2016) pp. 195–221.

14 S. Pressé, K. Ghosh, J. Lee, and K. A. Dill, Rev. Mod. Phys. 85, 1115 (2013).

15 P. D. Dixit, A. Jain, G. Stock, and K. A. Dill, J. Chem. Theor. Comp. 11, 5464 (2015).

16 J. M. L. Ribeiro and P. Tiwary, bioRxiv, 400002 (2018).

17 J. M. L. Ribeiro, P. Bravo, Y. Wang, and P. Tiwary, J. Chem. Phys. 149, 072301 (2018).

18 P. Tiwary and M. Parrinello, Phys. Rev. Lett. 111, 230602 (2013).

19 P. Tiwary, V. Limongelli, M. Salvalaglio, and M. Parrinello, Proc. Natl. Acad. Sci. 112, E386 (2015).

20 Y. Wang, E. Papaleo, and K. Lindorff-Larsen, eLife 5, e17505 (2016).

21 Y. Wang, O. Valsson, P. Tiwary, M. Parrinello, and K. Lindorff-Larsen, J. Chem. Phys. 149, 072309 (2018).

22 J. Mondal, N. Ahalawat, S. Pandit, L. E. Kay, and P. Vallurupalli, PLoS computational biology 14, e1006180 (2018).

23 V. A. Feher, J. D. Durrant, A. T. Van Wart, and R. E. Amaro, Curr. Op. Struc. Bio. 25, 98 (2014).

24 V. A. Feher, E. P. Baldwin, and F. W. Dahlquist, Nat. Struc. Mol. Bio. 3, 516 (1996).

25 L. Zheng, M. Chen, and W. Yang, Proc. Natl. Acad. Sci. 105, 20227 (2008).

26 P. Tiwary and B. J. Berne, J. Chem. Phys. 147, 152701 (2017).

27 J. Pfaendtner and M. Bonomi, J. Chem. Theor. Comp. 11, 5062 (2015).

28 O. Valsson and M. Parrinello, Phys. Rev. Lett. 113, 090601 (2014).

29 P. Tiwary and M. Parrinello, J. Phys. Chem. B 119, 736 (2014).

30 R. A. Copeland, Nat. Rev. Drug. Discov. 15, 87 (2016).

31 A. Dickson, P. Tiwary, and H. Vashisth, Curr. Top. Med. Chem. (2017).

32 A. C. Pan, D. W. Borhani, R. O. Dror, and D. E. Shaw, Drug Discov. Today 18, 667 (2013).

33 J. M. L. Ribeiro, S.-T. Tsai, D. Pramanik, Y. Wang, and P. Tiwary, arXiv preprint arXiv:1809.04540 (2018).

34 S. D. Lotz and A. Dickson, J. Amer. Chem. Soc. 140, 618 (2018).

35 P. Tiwary, J. Mondal, and B. J. Berne, Science Advances 3 (2017).

36 P. Tiwary, J. Phys. Chem. B 121, 10841 (2017).

37 K. L. Fleming, P. Tiwary, and J. Pfaendtner, J. Phys. Chem. A 120, 299 (2016).

38 C. D. Fu, L. F. L. Oliveira, and J. Pfaendtner, J. Chem. Phys. 146, 014108 (2017).

39 R. Casasnovas, V. Limongelli, P. Tiwary, P. Carloni, and M. Parrinello, J. Amer. Chem. Soc. 139, 4780 (2017).

40 M. Salvalaglio, P. Tiwary, and M. Parrinello, J. Chem. Theor. Comp. 10, 1420 (2014).

41 S. Kirkpatrick, C. D. Gelatt, and M. P. Vecchi, Science 220, 671 (1983).

42 A. Barducci, G. Bussi, and M. Parrinello, Phys. Rev. Lett. 100, 020603 (2008).

43 B. Hess, C. Kutzner, D. Van Der Spoel, and E. Lindahl, J. Chem. Theor. Comp. 4, 435 (2008).

44 G. A. Tribello, M. Bonomi, D. Branduardi, C. Camilloni, and G. Bussi, Comp. Phys. Comm. 185, 604 (2014).

